# Diverse Pathophysiological Processes Converge on Network Disruption in Mania

**DOI:** 10.1101/430868

**Authors:** Ivy Lee, Kathryn Nielsen, Mei-Hua Hall, Dost Öngür, Matcheri Keshavan, Roscoe Brady

**Affiliations:** Department of Psychiatry, Beth Israel Deaconess Medical Center, Boston, MA, USA.; Department of Psychiatry, Harvard Medical School, Boston, MA, USA.; Schizophrenia and Bipolar Disorder Program, McLean Hospital, Belmont, MA, USA.

**Keywords:** fMRI, lesion, mania, network, bipolar, disorder

## Abstract

**Background:** Neuroimaging of psychiatric disease is challenged by the difficulty of establishing the causal role of neuroimaging abnormalities. Lesions that cause mania present a unique opportunity to understand how brain network disruption may cause mania in both lesions and in bipolar disorder.

**Methods:** A literature search revealed 23 case reports with imaged lesions that caused mania in patients without history of bipolar disorder. We traced these lesions and examined resting-state functional Magnetic Resonance Imaging (rsfMRI) connectivity to these lesions and control lesions to find networks that would be disrupted specifically by mania-causing lesions. The results were then used as regions-of-interest to examine rsfMRI connectivity in patients with bipolar disorder (n=16) who underwent imaging longitudinally across states of both mania and euthymia alongside a cohort of healthy participants scanned longitudinally. We then sought to replicate these results in independent cohorts of manic (n=26) and euthymic (n=21) participants with bipolar disorder.

**Results:** Mania-inducing lesions overlap significantly in network connectivity. Mania-causing lesions selectively disrupt networks that include orbitofrontal cortex, dorsolateral prefrontal cortex, and temporal lobes. In bipolar disorder, the manic state was reflected in strong, significant, and specific disruption in network communication between these regions and regions implicated in bipolar pathophysiology: the amygdala and ventro-lateral prefrontal cortex.

**Limitations:** The was heterogeneity in the clinical characterization of mania causing lesions.

**Conclusions:** Lesions causing mania demonstrate shared and specific network disruptions. These disruptions are also observed in bipolar mania and suggest a convergence of multiple disorders on shared circuit dysfunction to cause mania.

## 1. Introduction

Bipolar disorders are common and severe but their pathophysiology is poorly understood and existing treatments are only partially effective with significant side effects(Chen and Dilsaver, 1996; Kessler et al., 1999; Leverich et al., 2003; Perlis et al., 2006; Slama et al., 2004). A wealth of studies have used functional magnetic resonance imaging (fMRI) in humans to understand bipolar pathophysiology at the network and circuit level. Fewer studies, however, have compared bipolar participants in different symptomatic states. In the clinic, a common goal of treatment is to shift individuals from a pathologic, symptomatic state (e.g. mania or depression) to an asymptomatic state of euthymia. Without a biological mechanism that accounts for these different states, it is unclear what process we should be targeting in the development and evaluation of therapeutics or what animal models accurately recapitulate bipolar disease pathophysiology.

Our group and others have utilized fMRI to try and understand how different bipolar mood states are reflected in brain network activity and connectivity. Previous studies utilized task-based fMRI to contribute to a model of how bipolar mood state is reflected in brain activation during emotion processing and regulation (Almeida et al., 2010; Cerullo et al., 2012; Chen et al., 2006; Chen et al., 2010; Hummer et al., 2013; Liu et al., 2012; Perlman et al., 2012; Rive et al., 2015; Strakowski et al., 2016; Van der Schot et al., 2010; Versace et al., 2010). Complementary experiments have used task-free (‘resting-state’, rsfMRI) imaging to generate connectome-wide maps of altered brain communication in different mood states albeit without specific correlates to cognitive processes (Altinay et al., 2016; Anand et al., 2009; Brady et al., 2016; Brady et al., 2017b; Li et al., 2015; Spielberg et al., 2016).

These studies have provided a greater understanding of how mood state is reflected in brain activity and connectivity. A challenge to the interpretation of most neuroimaging studies (including our own) is the difficulty in establishing causal relationships between fMRI signals and psychopathology (Etkin, 2018). Significant differences in brain activation and connectivity can be found between bipolar mood states but these findings may reflect disease process, or the brain’s compensatory response to illness, or may be epiphenomena.

A way to address this problem would be to limit the neuroimaging hypotheses tested to those with preexisting evidence of causality. Specifically, existing literature supports the idea that acute disruption of brain activity in circumscribed regions can cause mania. If these injuries impinge on a shared brain substrate, we can generate of a model of how brain pathophysiology causes mania. This model could then be tested in bipolar disorder to determine if bipolar mania demonstrates the same relationship between behavior and pathophysiology.

Mania after an acute onset, localized brain lesion is rare, but well-documented in many case reports (reviewed in (Santos et al., 2011; Satzer and Bond, 2016)). Starkstein and Robinson used brain imaging to localize lesions in patients experiencing mania associated with a brain lesion e.g. (Robinson et al., 1988; Starkstein et al., 1988; Starkstein et al., 1990; Starkstein et al., 1987). They found these lesions were anatomically distributed, which raises an intriguing possibility: if these lesions did not converge on a unitary anatomical vulnerability to mania, maybe they shared a different common feature: *disruption of a shared brain network*. Starkstein et al. postulated such a model (Starkstein et al., 1988) but neuroimaging at the time did not allow visualization of network communication between regions. The relevance of these lesion induced manic syndromes to bipolar mania remained undetermined.

The hypothesis that disparate lesions may cause a common phenotype via disruption of a shared brain network is well validated (Carrera and Tononi, 2014; Feeney and Baron, 1986; He et al., 2007; Honey and Sporns, 2008). The broader application of this understanding has been limited by the challenges of 1) performing imaging in acutely ill clinical populations and 2) accumulating a sufficient number of cases to make valid inferences about the generalizability of results. A recent innovation to circumvent these challenges is an imaging analysis method termed “lesion network mapping” (Boes et al., 2015). This approach utilizes published cases of lesions associated with a behavioral phenotype of interest and, using brain connectivity data from healthy participants, demonstrates which network(s) the lesions would be expected to disrupt. Drawing on the historical record of these cases that both document the behavioral phenotype and include imaging has allowed sufficient sample size to discern behavior-specific patterns of network disruption in a variety of phenotypes. This approach has already been used to analyze a variety of phenotypes such as criminality (Darby et al., 2018), hallucinations (Boes et al., 2015), movement disorders (Fasano et al., 2017; Laganiere et al., 2016), and delusions (Darby et al., 2017).

We used lesion network analysis to determine if anatomically distinct lesions that cause mania converge on a common brain network(s). We then tested the generalizability of the relationship between network disruption and the mania phenotype by determining if networks disrupted in lesion-induced mania were also disrupted in patients with bipolar disorder mania. Our hypothesis was that diverse pathophysiological processes (e.g. ischemic strokes vs. bipolar disorder) can give rise to a shared behavioral phenotype of mania by disrupting the same brain network.

## 2. Materials and Methods

### 2.1 Methods Overview

In the first step of the analysis, we used lesion network mapping to determine if lesions that cause mania converge on a shared network. In the second step of our analysis, we tested the hypothesis that different pathophysiological causes of mania disrupt the same brain network by asking whether the disruption we observed in our lesion network mapping experiment was also present in bipolar disorder mania. This test was performed in a cohort of participants with bipolar disorder imaged longitudinally in both a manic state and a euthymic state to isolate state related changes in brain connectivity. We then tested this network in a replication dataset of two additional cohorts of bipolar participants.

#### 2.2.1 Lesion Network Mapping Overview

Lesion network mapping was performed using methods similar to Boes et al. (Boes et al., 2015). In brief, this involves three steps: First, we searched the literature for case reports of brain lesions causing mania. Next these lesions were traced by hand from MRI or CT scans onto a standardized template brain. We used each traced lesion as a separate region of interest (ROI) in a rsfMRI analysis using data from 40 healthy control participants. Finally, we compared the connectivity maps of lesions causing mania to the connectivity maps of randomly selected, publicly available brain lesion tracings (Liew et al., 2018). Brain regions that were significantly more likely to be functionally connected to mania inducing lesions are then reported.

#### 2.2.2 Case Reports of Lesion Induced Mania with Neuroimaging

We conducted a literature review to find cases of lesion-induced mania using a PubMed search with terms such as ‘mania AND lesion’ followed by searches of other literature referenced in those publications. Inclusion criteria were: 1) cases had to describe a temporally linked onset of neurological symptoms and mania suggesting a causal relationship between brain injury and manic symptoms. 2) Cases had to include published imaging (e.g. MRI or CT scan) that allowed tracing. 3) Patients could not have a pre-existing bipolar or psychotic disorder. 23 cases met these criteria, including many found in a recent review of stroke related mania (Santos et al., 2011). Case reports and references to original literature are included in Appendix Table 1. The MRI and CT images included with these published case reports were used as guides to hand trace lesions onto a standardized Montreal Neurological Institute 152 (MNI152) template brain using FSL (https://fsl.fmrib.ox.ac.uk/fsl/fslwiki/) and 3D Slicer (https://www.slicer.org/). (Figure 1) Lesion tracing fidelity to the published image was confirmed by consensus agreement between I.L. and R.B.

**Figure 1:**
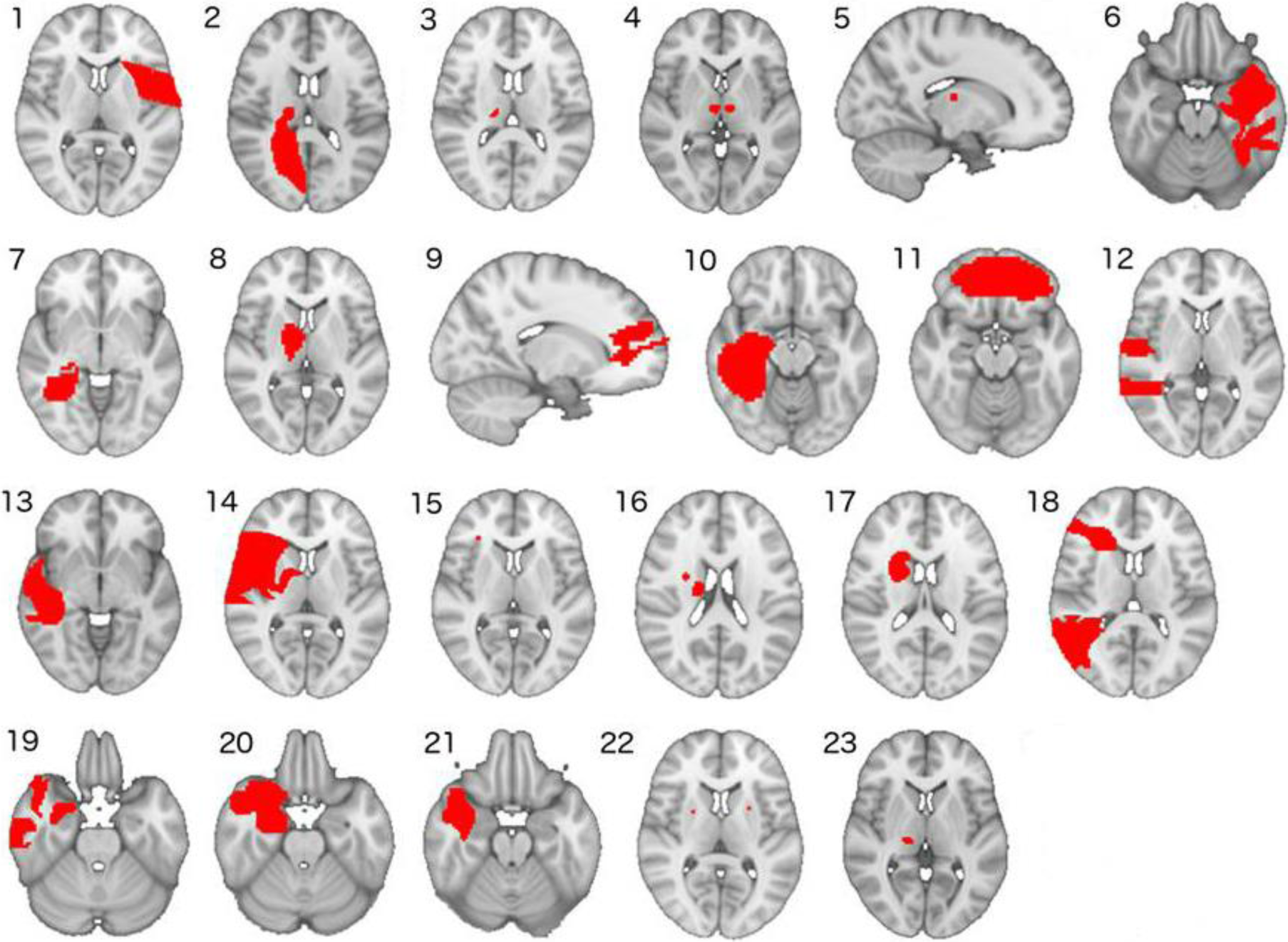
Tracings of Lesions that Induced Mania. Lesion location of twenty-three cases of lesion-induced mania, here traced onto a standard template brain. Horizontal sections are oriented with the right side of the brain on the reader’s left.

#### 2.2.3 Normative resting-state fMRI data for lesion mapping

rsfMRI data was acquired from a cohort of 40 healthy control (HC) participants (20 female, 20 male. Age: mean 29.4, SD +/−10.7). Details of their assessment are included in Supplemental Methods. All MRI data was acquired on a 3T Siemens Trio-TIM scanner using a standard 12-channel head coil. No scanner upgrade occurred during the duration of data collection. Scanner sequences and subsequent imaging data preprocessing steps were the same as performed in previous studies and are detailed in Supplemental Methods (Brady et al., 2017a; Brady et al., 2016; Brady et al., 2017b). Preprocessing methods were chosen to minimize motion artifacts.

#### 2.2.6 Connectivity Maps to Mania Inducing Lesions

The time course of voxels in each traced lesion ROI was extracted and Pearson correlation coefficients between this time course and those of all other voxels in all HC participants were calculated using DPABI (Yan et al., 2016). These values were transformed to Fisher’s z scores to generate FC maps of regions functionally connected to mania inducing lesion ROIs. One-sample t-tests of these maps from all 40 participants were generated using SPM. Resulting connectivity maps were thresholded at a p value <.0001. The resulting maps of functionally connected areas were binarized using FSL (Figure 2).

**Figure 2:**
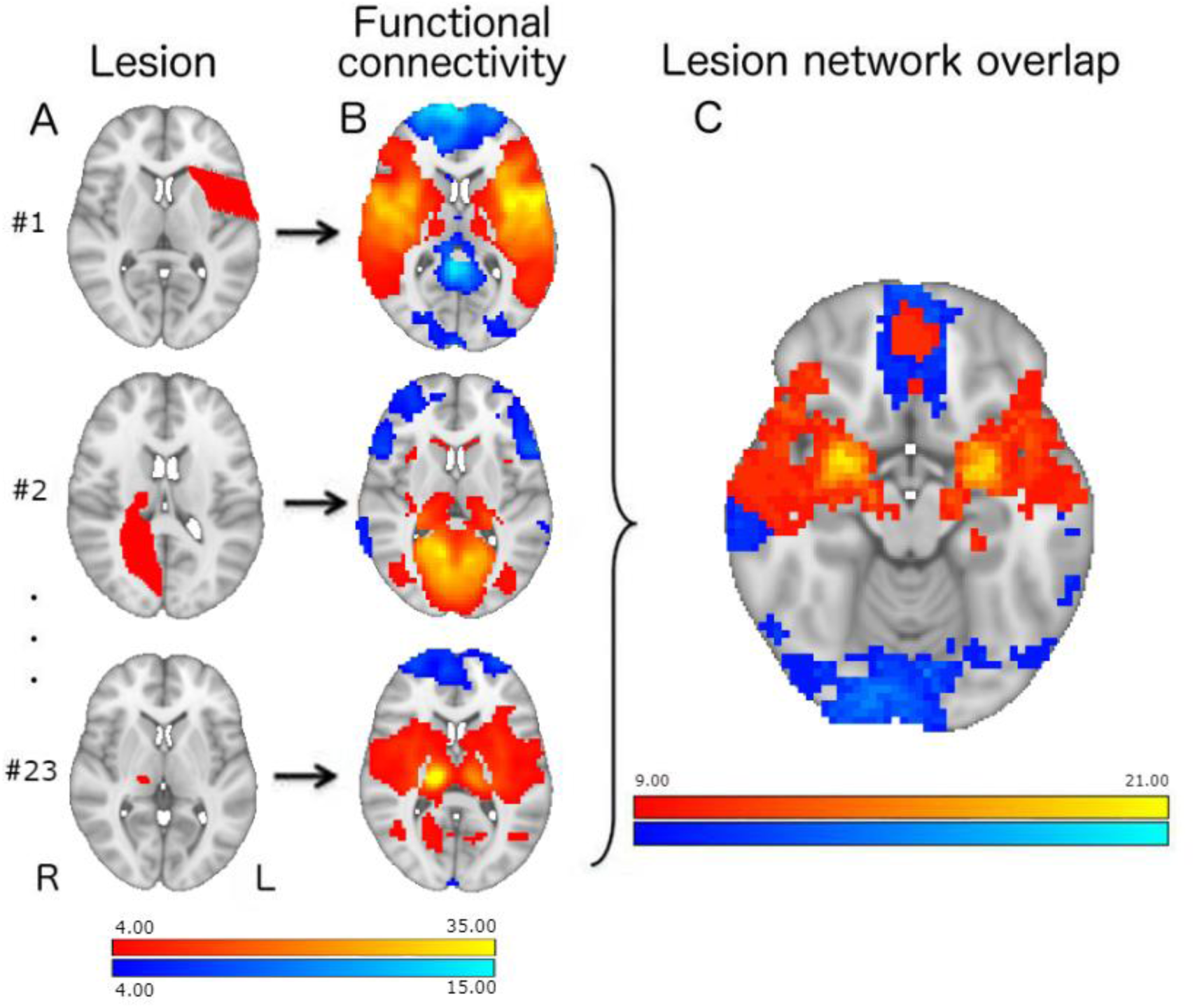
Network Mapping for Mania Inducing Lesions. Twenty-three lesions that caused mania were traced from case reports with published images onto a template brain (column A). Each lesion tracing was used as a ROI for mapping functional connectivity to this region in healthy control subjects (B), allowing visualization of the network that would be disrupted by the lesion. Regions of positively correlated activity to the ROI are shown in hot colors and negatively correlated activity in cold colors. These lesion network maps were overlaid to identify convergent areas of network connectivity shared among mania inducing lesions (C). Left Color Bar: T-statistic. Right Color Bar: Number of lesion maps converging on a given voxel.

#### 2.2.7 Comparison with Random Control Brain Lesions

Mania after a brain lesion is a rare outcome. For disruption of network connectivity to be a plausible explanation for this behavioral phenotype, these lesions would have to disrupt connectivity in networks not normally disrupted in most brain lesions. To control for lesion specificity, we made use of a publicly available database of hand traced stroke lesions (Liew et al., 2018). We randomly selected 23 lesions from the database and completed the same process described above under ‘Connectivity Maps to Mania Inducing Lesions’ to generate maps of connectivity to control brain lesions. As in prior lesion network mapping studies, we compared the connectivity maps of mania-inducing lesions to maps generated from control lesions using a voxel-wise Liebermeister test (Rorden et al., 2007). This test generates a map of voxels that are more likely to be connected to mania inducing lesions than control lesions. Multiple comparison correction in the Liebermeister test occurs via False Discovery Rate (FDR) correction.

#### 2.2.8 Comparison with Size Matched Control Brain Lesions

As an additional control for the network specificity of our mania causing lesions, we compared the 23 mania causing lesions to another set of entirely distinct 23 control brain lesions that were chosen on the basis of size matching to the mania causing lesions. Connectivity maps to this second set of lesions were again compared to the networks derived from mania causing lesions using the same Liebermeister test described above.

#### 2.2.9 Comparison with Single-Plane Traced Control Brain Lesions

Lesions in lesion network analysis are traced in a single plane by virtue of data presented (i.e. single plane images in case report figures). Prior studies that directly compared representative single-plane and 3-D tracings saw high concordance (spatial correlation coefficient of 0.96) in the resultant network connectivity maps (Boes et al., 2015). While this method has been accepted in all prior lesion network mapping studies, we opted to be conservative and compared the connectivity maps from the single-plane mania-causing lesion tracings to a set of 23 control lesions that were re-traced in a single plane (i.e. a direct comparison of mania-causing lesions traced in a single plane to control lesions traced in a single plane). Resultant connectivity maps were again compared using the Liebermeister test.

### 2.3 Comparison to Bipolar Disorder Mania Overview

We sought to compare the results of our mapping analysis from lesion induced mania to patients with mania resulting from bipolar disorder. We hypothesized that mania caused by either neurological lesions or bipolar disorder results from a shared pattern of network disruption. To test this, we collected rsfMRI data from a cohort of participants (n=16) with bipolar disorder type I who underwent longitudinal imaging during a manic state and then again during a euthymic state. We then used the results of the lesion mapping analysis above as ROIs to determine whether disruptions in similar networks were observed in bipolar mania (when compared to euthymia). We then sought to replicate this analysis an independent cohort of participants with bipolar disorder, comparing participants with bipolar disorder type I imaged while manic (n=26) to those imaged while euthymic (n=21).

#### 2.3.1 Bipolar & Healthy Comparison Longitudinal Participants

Study recruitment was performed as in our previous studies and as detailed in Supplemental Methods (Brady et al., 2017a; Brady et al., 2016; Brady et al., 2017b). Longitudinal participants with bipolar disorder type I were all recruited, characterized, and imaged while in a manic state. Participants were then contacted subsequently for imaging and clinical characterization while euthymic. Of the cohort of bipolar participants with usable scan data at both time points, almost all (15/16) of were initially scanned while hospitalized on inpatient units in a manic state. All euthymic scans took place during outpatient treatment.

We also recruited a cohort of sex and age matched healthy control participants to scan longitudinally at a similar interval to the bipolar cohort in order to control for non-specific effects such as scanner drift.

#### 2.3.2 Bipolar Replication Dataset Participants

For the replication dataset, we recruited participants with bipolar disorder type I that were recruited, characterized, and imaged a single time each during either a manic state (n=26) or a euthymic state (n=21). Recruitment and characterization was otherwise performed identically to the longitudinal participants described above.

#### 2.3.3 MRI data Acquisition & fMRI data processing

Data acquisition for bipolar longitudinal imaging was collected on the same scanner using the same sequence as above. fMRI data processing was performed exactly as described above. No scanner upgrades occurred during the duration of the study.

#### 2.3.4 Bipolar Mania versus Euthymia Resting-State Longitudinal Connectivity Analysis

Regions identified in the lesion mapping study were used as ROIs to examine whole-brain connectivity differences using a within-subject design. The time course of voxels in each ROI was extracted and Pearson correlation coefficients between this timecourse and those of all other voxels were calculated using DPABI (Yan et al., 2016). These values were transformed to Fisher’s z scores to generate FC maps of regions functionally connected to these ROIs. Maps were then entered into a second level design in SPM using a paired t-test comparing the manic state to the euthymic state within-subjects. Given concerns about false positives resulting from more permissive voxel thresholds (Eklund et al., 2016), plus concerns about the reproducibility of spatially constricted, highly thresholded imaging signals (Duncan et al., 2009), we required that the resulting maps of mood state differences meet two different standards for statistical significance: First, thresholding at the voxelwise level of p<.001 with an extent threshold of k=28 voxels for p<.016 clusterwise significance (alpha <.05 corrected for three comparisons). Second, a voxelwise threshold of p<.005, k=59 voxels for p<.016 clusterwise significance. The threshold for cluster level significance was determined using Monte Carlo simulation as implemented in DPABI (Yan et al., 2016).

Regions of significant difference between bipolar mood states were used to generate ROI to ROI z-transformed functional connectivity values in the bipolar participants and in a cohort of healthy comparison (HC) participants imaged longitudinally. To control for non-specific effects (e.g. scanner drift), these results were entered into a two-way mixed ANOVA to determine the effect of time/mood state on diagnosis (bipolar vs. HC).

To control for within-subject differences in medication regimen we performed a linear mixed model analysis using SPSS statistical software (SPSS Inc., Chicago, IL, USA) to examine the effect of mood state with medications as time-varying covariates.

#### 2.3.5 Bipolar Mania versus Euthymia Resting-State Replication Cohort Connectivity Analysis

Regions identified in the lesion mapping study were used as ROIs to examine functional connectivity differences using a between-group design. The time course of voxels in each ROI was extracted and Pearson correlation coefficients between this timecourse and those of all other voxels were calculated using DPABI (Yan et al., 2016). These values were transformed to Fisher’s z scores to generate FC maps of regions functionally connected to these ROIs. Maps were then entered into a second level design in SPM using a independent sample t-test comparing the manic group to the euthymic group. Covariates included age, sex, and prescribed antipsychotic dosage (CPZE). Between-group connectivity analyses were confined to regions that were identified in the longitudinal bipolar cohort. Within these regions, voxelwise significance was thresholded at p<.05 and cluster extent thresholded significance set at p<.025 clusterwise significance (alpha <.05 corrected for two comparisons).

## 3. Results

### 3.1.1 Lesions temporally associated with mania are spatially heterogeneous

We found 23 cases of lesion induced mania with imaging data (Appendix Table 1). In all cases these patients had no prior history of bipolar or psychotic disorder and only became manic following the development of a lesion. The 23 lesions were spatially diverse (Figure 1). When overlaid the highest area of overlap was between 3 lesions in the thalamus.

### 3.1.2 Lesions that cause mania show substantial overlap in functional connectivity

In contrast to the heterogeneity in location observed among mania causing lesions, these lesions demonstrated substantial overlap in functional connectivity. This was observed in the salience network (Dosenbach et al., 2008; Seeley et al., 2007) broadly and most prominently in the right amygdala (20/23 lesions). Given that the amygdala is a broadly connected network ‘hub’ (van den Heuvel and Sporns, 2013) this is not surprising. Indeed, in the set of 23 randomly selected control stroke tracings these lesions were commonly functionally connected to the bilateral amygdala (15/23 lesions).

### 3.1.3 Mania causing lesions are functionally connected to regions that are not connected to control lesions

When the connectivity maps to mania lesions were compared to maps from control lesions, three brain regions were significantly more likely to be functionally connected to mania causing lesions than control lesions: 1) The bilateral temporal lobe, most prominently in the temporal poles but extending to the hippocampus. 2) The medial orbitofrontal cortex (OFC), and 3) Brodmann’s area 46 (Dorso-Lateral Prefrontal Cortex, DLPFC) (Figure 3). The temporal regions and OFC were correlated with mania causing lesions and the BA46 region was anti-correlated with mania causing lesions. Notably, the areas of significant differences were *only* functionally connected to mania inducing regions i.e. none of the 23 random control lesions were functionally connected to these regions. We continued to observe the same pattern of differential connectivity to mania-causing lesions under multiple control comparisons i.e. when compared to random ischemic lesions, a distinct set of size matched ischemic lesions, and ischemic lesions traced in a single plane (Supplemental Figures 1, and 2).

**Figure 3:**
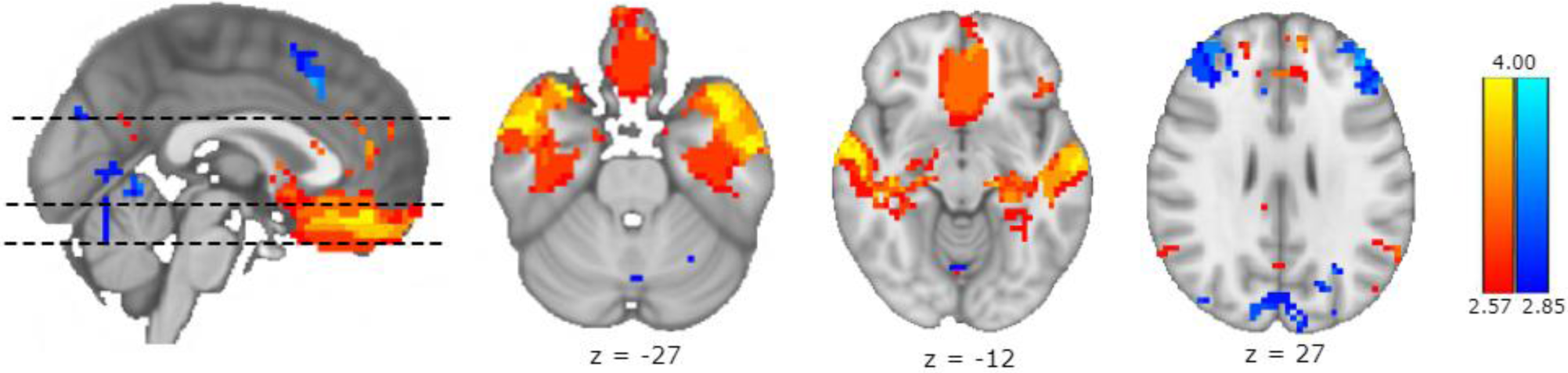
Anatomical Specificity of Lesion Induced Mania. Network connectivity to mania causing lesions is compared to network connectivity in a set of control stroke lesions. A voxelwise Liebermeister test reveals regions significantly (p-FDR <.05) more functionally connected to mania-causing lesions than controls lesions. Regions of positively correlated activity to the ROI are shown in hot colors and negatively correlated activity in cold colors. Three regions of significant differences in connectivity were observed: The Orbitofrontal Cortex (OFC), the bilateral temporal lobe, and the bilateral Dorso-Lateral Pre-Frontal Cortex (DLPFC, Brodmann Area 46). The OFC and temporal lobes were intrinsically correlated with areas of mania-causing lesions. BA46 was intrinsically anti-correlated with mania-causing lesions. None of the control lesions were connected to these areas. Color Bar: Z-stat.

As an exploratory analysis, we asked if these network connectivity relationships could explain *all* cases of lesion-induced mania. Fully 12 of 17 non-thalamic lesions were in locations functionally connected to the regions described above (i.e. regions that were not connected to *any* of the random control lesions). The 6 thalamic lesions that caused mania were all located in the right hemisphere and functionally connected to the salience network.

We also performed a post-hoc analysis to reduce heterogeneity among the pathological mechanisms involved in mania causing lesions. When we confined our lesions of interest to those resulting from an infarct (n=13) and compared these network maps to the 23 control lesions, we continued to observe the same mania lesion specific connectivity in the temporal lobes and orbito-frontal cortex but the previously observed region of anti-correlation to mania causing lesions in BA46 was no longer statistically significant (Supplemental Figure 3).

### 3.2 Extension of Lesion Induced Mania Studies to Bipolar Disorder Mania

#### 3.2.1 Bipolar Disorder Longitudinally Scanned Participants

Bipolar participants were significantly more symptomatic on all measures (YMRS, MADRS, PANSS, PANSS positive sub-score) during the manic state than during the euthymic state (Supplemental Table 2). Increased MADRS scores in manic subjects typically came from scores on the reduced sleep, concentration difficulties, and inner tension items. Symptom scales were not available for one participant who was euthymic on SCID. Medication regimens did not differ significantly between these two time points.

#### 3.2.2 Networks Disrupted in Lesion Induced Mania are Disrupted in Bipolar Disorder Mania

We compared within-subject differences in functional connectivity between bipolar mania and euthymia in the three regions (temporal poles, OFC, and BA46) identified in lesion-induced mania above. Consistent with our original hypothesis we did observe significant and specific disruptions in connectivity when comparing bipolar mania to euthymia (Figure 4): The temporal lobes ROI demonstrated decreased connectivity to the bilateral ventro-lateral prefrontal cortex (VLPFC) in mania compared to euthymia. No significant increases in connectivity to the temporal poles ROI in mania were observed. The BA46 ROI demonstrated increased connectivity to the left amygdala and dorso-medial prefrontal cortex in mania compared to euthymia. No significant decreases in connectivity to the BA46 ROI in mania were observed. Of note, BA46 is normally anti-correlated to the amygdala such that this result and the temporal pole-VLPFC are all consistent with a disruption of coordinated (i.e. either correlated or anti-correlated) activity in mania when compared to euthymia. The orbitofrontal ROI did not demonstrate significant differences in functional connectivity when comparing mania to euthymia.

**Figure 4:**
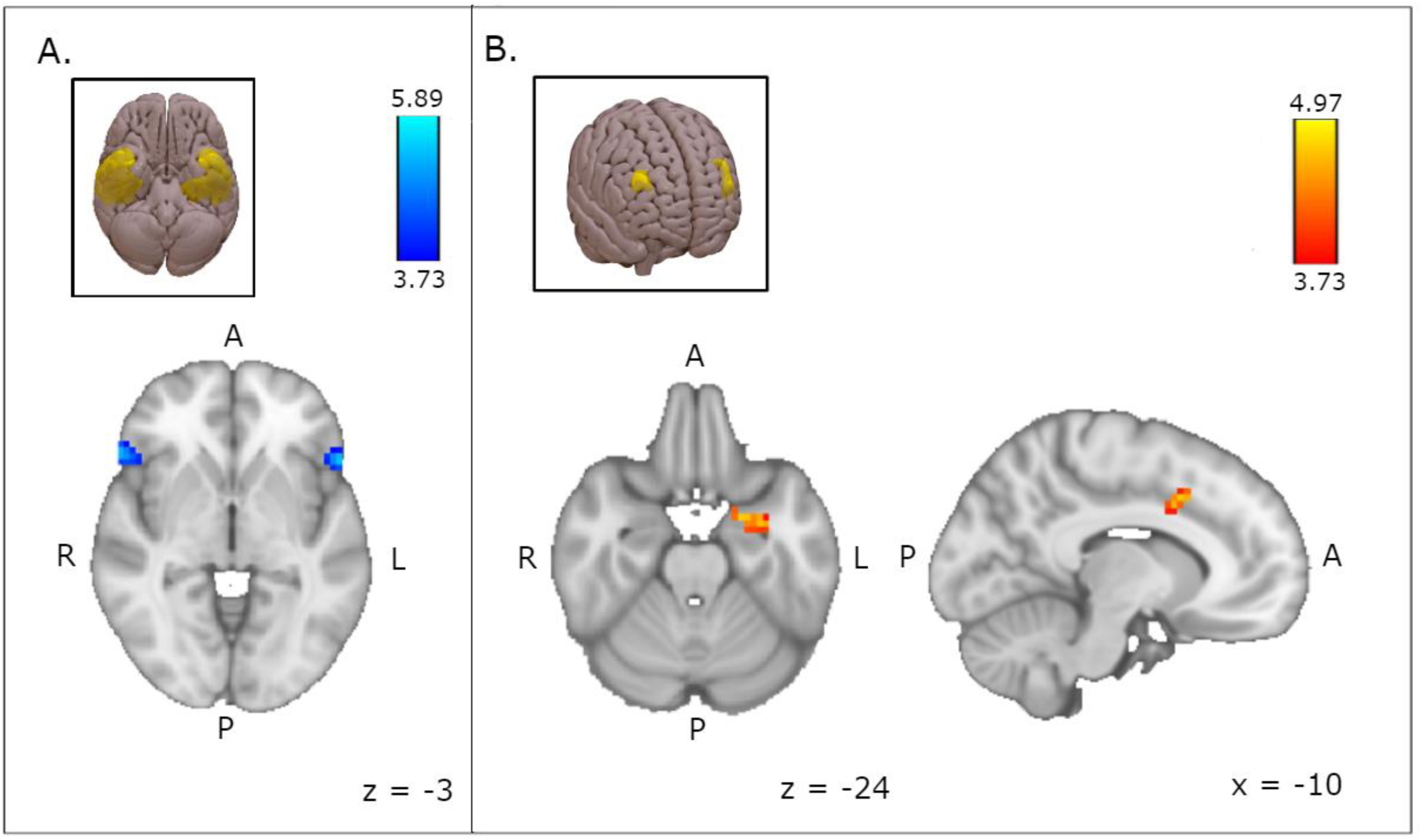
Lesion Induced and Bipolar Mania Converge On Shared Network Dysconnectivity. We sought to determine if the networks identified in the lesion mapping analysis of mania also disrupted in bipolar mania. Using the regions identified in lesion induced mania as ROIs, we compared functional connectivity to these areas in participants with Bipolar Disorder type I imaged in a manic state and then in a euthymic state. A) Using the temporal pole ROIs from lesion network mapping (inset), we observe disconnectivity in mania to the bilateral Ventro-Lateral Pre-Frontal Cortex (VLPFC) (Right: k=47, peak voxel T-stat 5.89 p<.001, x+57 y+27 z0; Left: k=32, peak voxel T-stat 5.39 p<.001, x-54, y+21, z-3). B) The BA46 ROIs from the lesion mapping analysis (inset) demonstrate increased connectivity in mania to the left amygdala (k=29, peak voxel T-stat 4.74, p<.001, x-21 y0 z-24). Here increased connectivity means dis-connectivity between these normally anti-correlated structures. BA46 also demonstrated increased connectivity to the dorso-medial Pre-Frontal Cortex (dmPFC) (k=32, peak voxel T-stat 4.97, p<.001, x-12, y+9, z+36). A set of healthy comparison participants imaged longitudinally across these time points did not demonstrate any significant change in connectivity in these regions. Images are thresholded voxelwise p<.001, cluster-extent p<.016. Color Bar: T-stat.

Examining two connectivity relationships in more detail, the BA46-Left Amygdala ROIs connectivity showed a strong effect of mood state (T(1,15)= 6.236 p=.000016, Cohen’s d=1.56) with these regions effectively disconnected in mania (mean zFC -.022) compared to euthymia (mean zFC-.252). Temporal lobe-VLPFC connectivity also showed a strong effect of mood state (T(1,15)= 5.659, p=.000045, Cohen’s d=1.42) with these regions also effectively disconnected in mania (mean zFC -.003) compared to euthymia (mean zFC .277). See Supplemental Figure 4 for individual plots of connectivity in each participant in both time points.

Examining connectivity in these ROIs in both bipolar participants and longitudinally imaged HC participants to control for non-specific (e.g. scanner) effect, we continue to observe a statistically significant interaction : Using a mixed ANOVA with time as within-subject factor and diagnosis (bipolar vs HC) as a between group factor we continue to observe a significant time*state interaction in both the BA46-amygdala ROIs (F(1, 29) = 14.203, *p* = .0001, partial η^2^ = .329) as well as the temporal lobe-Right VLPFC ROIs (F(1, 29) = 14.734, *p* = .001, partial η^2^ = .337).

We did not observe any significant within-subject changes in medication regimen in the bipolar participants. Nevertheless, we re-performed these connectivity analyses with medication status as a time-varying covariate. These results remained highly significant (Supplemental Results). We did not exclude bipolar participants on the basis of a substance use disorder so we repeated these analyses excluding any participant that met criteria for a substance use disorder at either time point. This analysis continued to demonstrate significant effects of mood with similar or stronger effect sizes (Supplemental Results).

#### 3.2.2 Partial Replication of Bipolar Disorder Results in an Independent Cohort

Reproducibility in clinical neuroimaging studies is a significant concern. We sought to replicate our longitudinal bipolar imaging analysis in an independent cohort. While we did not have access to another cohort of bipolar patients with longitudinal imaging across mood states, we were able to compare two separate cohorts of bipolar disorder type I participants: One group imaged in a manic state (n=26) and the other imaged in a euthymic state (n=21) (Clinical / demographic information in Supplemental Table 3). We examined functional connectivity between the regions identified in the lesion network analysis and the regions identified in the longitudinal bipolar cohort. We observed the same pattern of disconnectivity between the temporal lobe ROI and the right VLPFC in the bipolar mania cohort when compared to the bipolar euthymia cohort. We did not observe the mood state dependent differences in BA46 connectivity that we observed in the longitudinal cohort.

## 4. Discussion

Here we observe that mania causing lesions do not show significant co-localization at the anatomic level but instead impinge upon shared brain networks. Two patterns of network localization are observed: For non-thalamic lesions, mania causing lesions differed from non-mania causing lesions via a pattern of connectivity whose distribution includes territories (OFC, BA46, temporal poles) that were never observed to be connected to locations of control lesions. All thalamic lesions that caused mania were functionally connected to the salience network suggesting that disruption of the thalamic node of this network may have different behavioral consequences than disruption of other nodes in this network, consistent with the recent discovery that the thalamus actually controls cortical functional connectivity (Schmitt et al., 2017).

We explored the generalizability of our findings by using the same fMRI imaging modality to determine if mania in bipolar disorder is reflected in disruption of the same networks. Analyzing data from bipolar participants scanned longitudinally across mood states we observe exactly this result. Specifically, we observed disconnectivity in mania within this network in structures previously implicated in the pathophysiology of bipolar disorder (i.e. amygdala and VLPFC) but heretofore not linked causally to mania (Blond et al., 2012; Chase and Phillips, 2016; de Zwarte et al., 2014; Womer et al., 2009)). Notably, most imaging studies of bipolar disorder that identify signals in these regions either investigate these regions as *a priori* regions of interest or use paradigms known to activate these structures (e.g. emotion processing tasks). We observe that completely distinct pathophysiological processes converge on both a shared behavioral phenotype and shared network disruption, particularly in structures long implicated in bipolar disorder pathophysiology but here identified empirically.

While this work draws upon a convergence of multiple pathological processes to identify a shared network basis for the emergence of mania, all of these analyses rely on a single imaging modality (rsfMRI connectivity). Other imaging modalities also implicate this network in bipolar disorder. Two recent studies examined structural connectivity and controllability (i.e. how neuronal activity propagates in local networks) in both bipolar participants and participants at genetic high risk for bipolar disorder (Jeganathan et al., 2018; Roberts et al., 2018). Using structural (not functional) MRI data, these analyses identified structural dysconnectivity and control deficits in high risk participants in a right-lateralized network involving dorsomedial and ventrolateral prefrontal cortex, the superior temporal pole, putamen, and caudate nucleus, which all correspond to the territories identified in our analysis.

## 5. Limitations

Methodological limitations of our study include: 1) In studying published case reports, we are limited by a reliance on external reports of symptomatology and characterization. Standardized reporting of symptom severity (e.g. Young Mania Rating Scales, SCID criteria for a manic episode) are not available in these cases or in other syndromes analyzed using lesion network mapping (Boes et al., 2015; Darby et al., 2018; Darby et al., 2017; Fasano et al., 2017; Laganiere et al., 2016). Despite this limitation, the cases analyzed demonstrate a significant network co-localization that also distinguishes them from randomly selected brain lesions and supports the hypothesis that these cases share a common biological feature of network co-localization. 2) We used HC rsfMRI data to estimate patterns of network connectivity that would be disrupted by lesions. The patients in the lesion induced mania studies were, on average, older than our HC dataset and therefore might have age-related differences in intrinsic connectivity from the HC dataset. We decided to use this HC dataset because it was collected on the same scanner and used the same scan sequence as the bipolar participant data. Furthermore, prior reports comparing lesion network mapping in young and elderly rsfMRI data did not give divergent results (Boes et al., 2015). 3) The control lesions we used for comparison to mania causing lesions were taken from an open resource that did not detail the post-stroke behavior of the patient. It is possible though exceedingly unlikely that these control lesions could have contained a lesion that caused mania. This would presumably cause a false negative result.

Limitations in imaging bipolar participants include: 1) For manic participants, motion confounds of rsfMRI data is an obvious concern. We attempted to minimize these through a combination of procedures that have been shown effective in removing spurious motion effects (Power et al., 2014; Yan et al., 2013). 2) The participants of our study were taking medications at the time of imaging. Our hypothesis that a within-subjects design would hopefully minimize medication changes between scans was proven correct, as we found no significant differences in regimens between mood states. The preponderance of evidence in bipolar neuroimaging suggests that the effects of medications normalize differences rather than exacerbating them (Hafeman et al., 2012). From both an ethical and a pragmatic standpoint, imaging participants while hospitalized for mania and un-medicated and then re-imaging while in a euthymic, un-medicated state appears untenable and unfeasible.

Finally, this study is fundamentally an observational study. We have not performed an intervention to demonstrate that network manipulation directly causes mania. That said, we have used cases where a brain injury appears causally linked to mania to constrain our hypotheses that we then test in bipolar disorder. Definitive testing the ability of any perturbation to cause a manic state in humans remains a challenge, but we believe the data presented here supports a causal role for network disruption in bipolar mania.

## 6. Conclusions

We hypothesize that both circumscribed anatomical lesions and a presumably pan-cellular disease can give rise to a shared manic phenotype via disruption of brain networks that mediate the behavioral phenotype. Specifically, this network can be disrupted by either acute injury at critical nodes (e.g. temporal lobe stroke) or by an interaction between environmental stressors and heritable pathology (e.g. altered signal transduction (Tobe et al., 2017) or excitability(Mertens et al., 2015)). (Figure 5)

**Figure 5:**
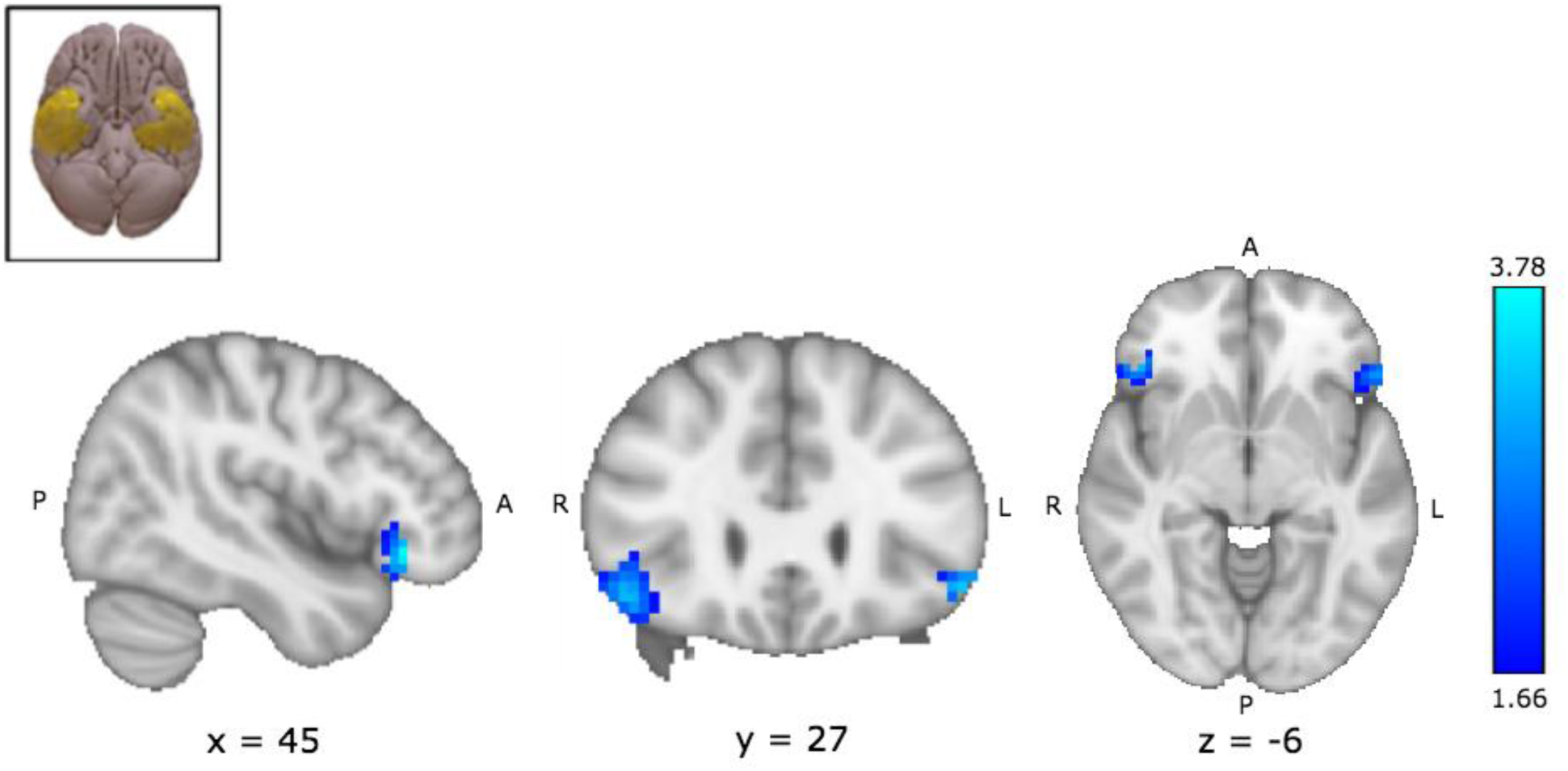
Replication of Temporal-VLPFC Disconnectivity in Bipolar Mania Compared to Euthymia. We sought to determine if the networks identified in the lesion mapping analysis of mania also disrupted in bipolar mania. Using the regions identified in lesion induced mania as ROIs, we compared functional connectivity to these areas in participants with Bipolar Disorder type I imaged in a manic state and then in a euthymic state. A) Using the temporal pole ROIs from lesion network mapping (inset), and examining the VLPFC regions identified in the longitudinal analysis (Figure 4), we again observe disconnectivity in mania to the bilateral Ventro-Lateral Pre-Frontal Cortex (VLPFC) (Right: k=52, peak voxel T-stat 3.78, p<.001, x45 y30 z-9; Left: k=23, peak voxel T-stat 3.33 p=.001, x-51, y27, z-9). While the VLPFC of both hemispheres showed reduced connectivity to the temporal lobe ROIs in mania, only the right VLPFC met extent significance (k=33 for p<.025 corrected for multiple comparisons). Images are thresholded voxelwise p<.05. Color Bar: T-stat.

**Figure 6:**
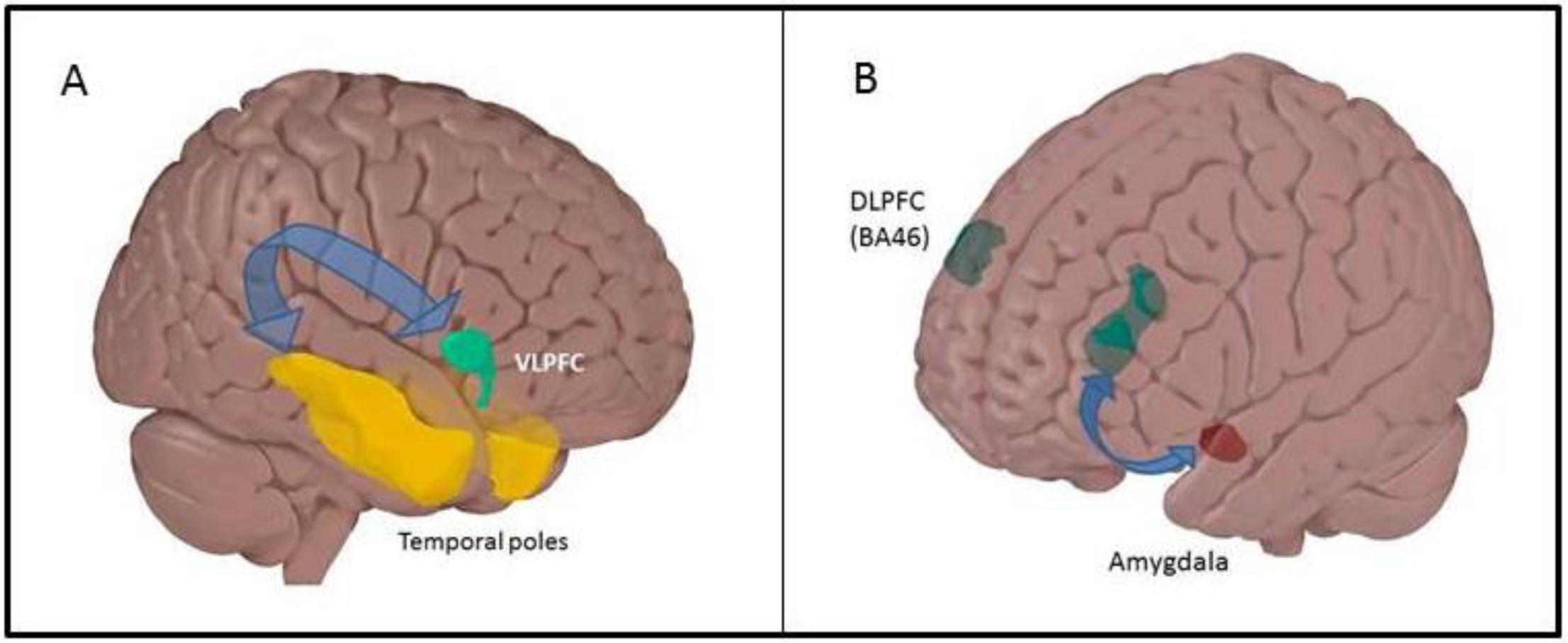
A Model of The Network Basis of Mania. We propose that multiple causes of mania converge on the same networks and it is disruption of distributed brain networks that mediate the link between pathophysiology and behavioral phenotype. Specifically, this network can be disrupted by either acute lesioning of critical nodes of this network (e.g. temporal lobes, OFC) or may be due to the combination of genetic factors plus environmental stressors that give rise to manic episodes in bipolar disorder. In either scenario, the disruption of communication across poly-synaptic networks causes the manic phenotype. Within these networks, data from longitudinal imaging supports disconnectivity between (A) Temporal poles and VLPFC and (B) Between amygdala and DLPFC (BA46) as causes of mania in bipolar disorder. Arrows here are not meant to denote a monosynaptic pathway.

Our findings support a hypothesis that disruption of this network is causally related to mania. Definitive testing of this hypothesis will require a non-human (e.g. primate) model system. One implication of our findings is that our approach may be generalizable to other symptoms and behaviors. The convergence of genetically determined and lesion-based causes of psychiatric symptoms on a finite set of circuit targets, provides an opportunity for a conceptual advance: Traditionally, animal models of psychiatric disorders are developed using perturbations of unclear relationships to disease pathology in humans (e.g. manipulation of candidate genes) followed by a search for plausible mouse behavioral analogues of human symptoms (e.g. motor activity). Our method of circuit identification may allow scientifically-based prospective laboratory models based on pathophysiology rather than behavioral output.

---

**Supplemental Table 1:** Case Reports of Lesion Induced Mania.

| Case # | Reference | Symptoms | Etiology | Location | Age |
| --- | --- | --- | --- | --- | --- |
| 1 | (Nagaratnam et al., 2006) (1) | Impulsivity, stubbornness, pressured speech, incoherency, aimless wandering, mood fluctuations, agitation, verbal aggression, euphoria, flight of ideas | Infarction | Left middle cerebral artery, frontal insula cortex, parietal regions | 72 |
| 2 | (Nagaratnam et al., 2006) (2) | Left sided weakness, blurred vision, mood fluctuation, hostility, hallucinations, insomnia, agitation, pressured speech, impulsivity | Infarction | Right posterior cerebral artery, right occipital lobe, thalamus | 80 |
| 3 | (Inzelberg et al., 2001) | Euphoric, talkative, spending a lot of money, jerky movements of his left limbs, distractible, pressured speech, flight of ideas, hypersexuality | Infarction | Thalamus, pulvinar | 61 |
| 4 | (Benke et al., 2002) | Vertical gaze paralysis, slight dysarthria, restlessness, agitation, euphoria, grandiosity, tangential and incoherent forms of speech | Infarction | Dorsalmedial thalami, intralaminar nuclei, interior nuclei | 38 |
| 5 | (Kulisevsky et al., 1993) | Euphoria, talkative, grandiose delusions, distractible, pressured speech, flight of ideas, decreased need for sleep, chorea, ballistic movements of left limbs | Infarction | Ventral lateral nuclei, ventromedial nuclei, thalamus | 81 |
| 6 | (Liu et al., 1996) | Sensory aphasia, depression, sleep disturbance, inappropriate guilt, poor appetite, ideas of death, irritability, hyperactivity, talkativeness, grandiose ideation | Infarction | Left temporal lobe | 48 |
| 7 | (Starkstein et al., 1988) (1) | Loud speech, inappropriate affect, change in affect, insomnia, irritability and suspiciousness | Infarction | Inferomedial temporal lobe | 66 |
| 8 | (Starkstein et al., 1988) (2) | Headaches, slurred speech, left sided weakness, euphoria, hyperactivity, pressured speech, flight of ideas, grandiose delusions, suspiciousness, hypersexuality, decreased need for sleep | Hemorrhage | Right thalamus | 36 |
| 9 | (Starkstein et al., 1988) (3) | Progressive blindness, insomnia, grandiose delusions, euphoria, hyperactivity, hypersexuality, talkative | Meningioma | Anterior inferior frontal region, crista galli, suprasellar cistern | 63 |
| 10 | (Starkstein et al., 1988)(4) | Left hemiparesis, left homonymous hemianopia, left hemihypesthesia, euphoria, hyperactivity, pressured speech, decrease need for sleep, headaches, left sided weakness | Meningioma | Temporal lobes, occipital lobes | 61 |
| 11 | (Starkstein et al., 1988) (5) | Hypersexuality, pressured speech, aggressive speech, irritability, threatening behavior, blindness in left eye, inability to converge eyes, tangential speech, euphoria | Meningioma | Frontal lobes | 48 |
| 12 | (Starkstein et al., 1988) (6) | Aggressiveness, impulsivity, intense headaches, euphoria, pressured speech, grandiose delusions, flight of ideas, and hyperactivity | Astrocytoma | Right temporal lobe, right lateral ventricle, shift of midline structures | 28 |
| 13 | (Rocha et al., 2008) | Irritability, emotional lability, decreased need for sleep, rapid speech and thoughts | Infarction | Right temporal-parietal lobe | 57 |
| 14 | (Mimura et al., 2005) | Depressed mood, psychomotor retardation, anxiety, irritability, delusions of guilt, euphoria, talkativeness, jocularity, grandiosity | Infarction | Middle cerebral arteries | 78 |
| 15 | (Starkstein et al., 1990) (1) | Hyperactivity, grandiose delusions, decreased need for sleep, pressured speech, mild irritability | Hemorrhagic contusion | Right frontal lobe | 37 |
| 16 | (Starkstein et al., 1990) (2) | Hyperactivity, euphoria, decreased need for sleep, pressured speech, grandiose ideation | Infarction | Ventral right internal capsule | 55 |
| 17 | (Starkstein et al., 1990) (3) | Decreased need for sleep, hyperactivity, euphoria, delusions, flight of ideas, hypersexuality, pressured speech | Infarction | Head of right caudate, anterior limb of internal capsule | 79 |
| 18 | (Starkstein et al., 1990) (4) | Unknown | Infarction | Basal temporal lobe, medial temporal gyrus, dorsolateral frontal area of cortex, head of the caudate | Unk. |
| 19 | (Starkstein et al., 1990) (5) | Unknown | Infarction | Basal temporal lobe, amygdala, hippocampus, head of the caudate | Unk. |
| 20 | (Starkstein et al., 1990) (6) | Unknown | Arteriovenous malformation | Right basal temporal lobe | Unk. |
| 21 | (Starkstein et al., 1990) (7) | Unknown | Hemorrhage | Right temporal lobe | Unk. |
| 22 | (Antelmi et al., 2014) | Left-sided limb weakness, insomnia, distractibility, increased energy, eccentric behavior | Lesion | Left caudate, bilateral putamen, left pons | 51 |
| 23 | (Julayanont et al., 2017) | Irritability, grandiosity, combative speech, paranoid delusion, high energy levels, decreased need for sleep, pressurized speech, euphoria, and inappropriate cheerfulness | Hemorrhage | Right thalamus, dorsomedial and pulvinar nuclei | 55 |

## Supplemental Methods

### Normative resting-state fMRI data for lesion mapping

rsfMRI data was acquired from a cohort of 40 healthy control (HC) participants (20 female, 20 male. Age: mean 29.4, SD +/−10.7) recruited as part of an ongoing study. All participants were recruited from the community as part of a protocol approved by the McLean Hospital Institutional Review Board, and all participants gave written informed consent before participating. Participants were assessed using the Structured Clinical Interview for the DSM-IV (SCID)(First and New York State Psychiatric Institute. Biometrics Research, 2007). Exclusion criteria were: Any current or historical psychiatric diagnosis, age outside the range of 18–65, any neurological illness, positive pregnancy test or lactation, history of head trauma with a significant loss of consciousness, and contraindications to magnetic resonance imaging.

### MRI data Acquisition

All MRI data were acquired on a 3T Siemens Trio-TIM scanner using a standard 12-channel head coil. No scanner upgrade occurred during the duration of data collection. Scanner sequences were the same as performed in previous studies and are detailed in Appendix Methods (Brady et al., 2017a; Brady et al., 2016; Brady et al., 2017b) Each scan consisted of two 6.2 minute rsfMRI runs with imaging parameters as follows: repetition time = 3000 milliseconds; echo time = 30 milliseconds; flip angle = 85°; 47 axial 3mm sections collected with interleaved acquisition. Structural data included a high-resolution, multiecho, T1-weighted, magnetization-prepared, gradient-echo image. All participants were told ‘remain still, stay awake, and keep your eyes open’. Video recording of the eyes was used to confirm the awake, eye-open state.

### fMRI data processing

Imaging data were preprocessed using Data Processing & Analysis of Brain Imaging (DPABI) (Yan et al., 2016). rsfMRI runs with head motion exceeding 3mm in any dimension or 3° of maximum rotation about three axes were discarded from further analysis. After realigning, slice timing correction, and co-registration, framewise displacement (FD) was calculated for all volumes (Power et al., 2012). Volumes with a FD greater than 0.2mm were regressed out during nuisance covariate regression. Any rsfMRI run with 50% or more of volumes regressed out was discarded from further analysis. Structural images from each subject were then normalized and segmented into gray, white and CSF partitions using the DARTEL technique(Ashburner, 2007). A Friston 24-parameter model was used to regress out head motion effects (Friston et al., 1996). The CSF and white matter signals, global mean signal as well as the linear trend were also regressed as nuisance covariates. The resultant data were band pass filtered to select low frequency (0.01–0.08Hz) signals. Functional images were brought into MNI (Montreal Neurological Institute) space and then smoothed by a Gaussian kernel of 8mm^3^ full-width at half maximum. Voxels within a gray matter mask were used for further analyses.

### Bipolar & Healthy Comparison Longitudinal Participant Recruitment and Characterization

Study recruitment was performed as in our previous studies (Brady et al., 2017a; Brady et al., 2016). All participants were recruited as part of a protocol approved by the McLean Hospital Institutional Review Board and gave written informed consent before participating. To ensure that participants understood the study, we conducted an informed consent survey, including simple questions about risks and benefits and the ability to withdraw consent. If the participants did not answer all questions correctly, the informed consent document was re-reviewed and understanding retested to ensure comprehension.

Longitudinal participants with bipolar disorder were all recruited, characterized, and imaged while in a manic state. Participants were then contacted subsequently for imaging and clinical characterization while euthymic. Of the cohort of bipolar participants with usable scan data at both time points, almost all (15/16) of were initially scanned while hospitalized on inpatient units in a manic state. All euthymic scans took place during outpatient treatment.

Diagnosis for all participants was determined using the Structured Clinical Interview for the DSM-IV (SCID)(First and New York State Psychiatric Institute. Biometrics Research, 2007)). All bipolar participants were also assessed using the Young Mania Rating Scale (YMRS), Montgomery–Asberg Depression Rating Scale (MADRS), and Positive And Negative Syndrome Scale (PANSS) at the time of imaging. Bipolar participants met criteria for the diagnosis of Bipolar Disorder Type I on both SCID and clinician report. A single participant diagnosed by multiple clinicians longitudinally as having bipolar disorder met SCID criteria for schizoaffective disorder. All participants had a history of psychosis during manic episodes. Bipolar participants scanned in a manic state met DSM-IV SCID criteria for a manic episode. One participant met criteria for a mixed episode. Participants imaged in a manic state also had a YMRS score of 20 or greater. Participants imaged in a euthymic state all did not meet DSM criteria for any mood episode for at least one month prior to imaging. Euthymic bipolar participants also had a YMRS score of 8 or less. All longitudinal HC comparison subjects had no current or historical diagnosis upon SCID assessment and no family history of first-degree relatives with psychiatric illness.

In the non-longitudinal bipolar cohorts, bipolar participants in the mania cohort were typically recruited from McLean Hospital inpatient units during hospitalization for a manic episode. Euthymic bipolar subjects were typically recruited from patients who had previously been hospitalized on these inpatient units. All replication cohort participants met SCID criteria for the diagnosis of bipolar disorder type I. Subjects imaged in a manic state all had a YMRS score of 20 or greater. Subjects in a euthymic state all had a YMRS score of 12 or less. There was no overlap in participants between the manic, euthymic, and longitudinal cohorts.

For both bipolar and healthy control participants exclusion criteria were: age outside the range of 18–65, neurological illness, pregnancy or lactation, electroconvulsive therapy in the last three months, history of head trauma with a loss of consciousness lasting more than a few minutes, and contraindications to magnetic resonance imaging.

### Bipolar Mood State Clinical / Demographic Analysis

For the longitudinal bipolar disorder participants, comparisons of the presence or absence of medications in subject’s regimens at both mood states used McNemar’s test for significant differences. Comparisons of symptom scales and the prescribed dosage of antipsychotic medications (chlorpromazine equivalents, CPZE) used a paired t-test.

## Supplemental Figures

**Supplemental Figure 1:**
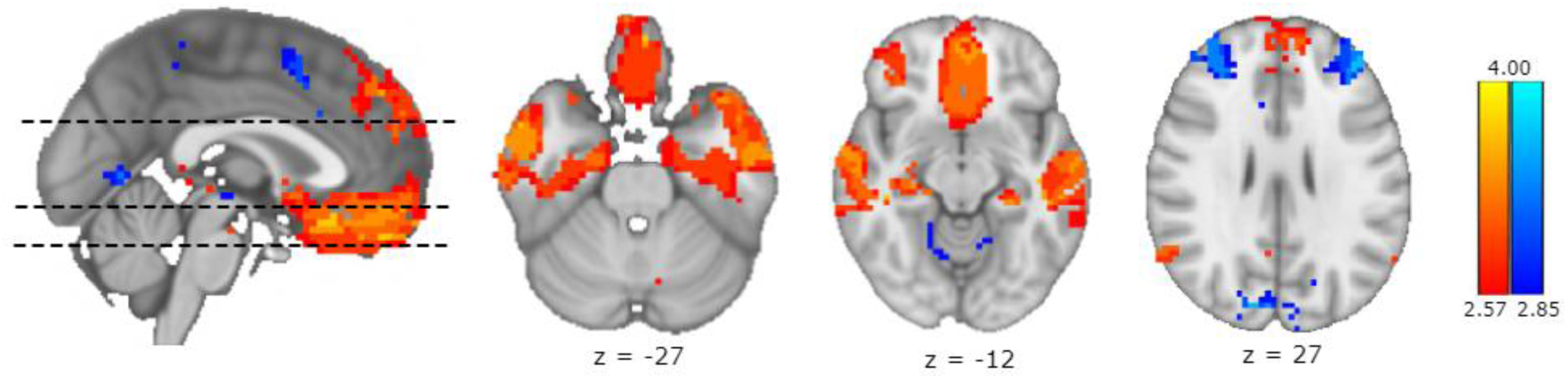
Mania Lesion Connectivity Compared to Size Matched Control Lesions: Network connectivity to mania causing lesions is compared to network connectivity in a set of control stroke lesions that have been selected on the basis of similarity in size to the mania causing lesions. A voxelwise Liebermeister test reveals regions significantly (p-FDR <.05) more functionally connected to mania-causing lesions than controls lesions. Regions of positively correlated activity to the ROI are shown in hot colors and negatively correlated activity in cold colors. Three regions of significant differences in connectivity were observed: The Orbitofrontal Cortex (OFC), the bilateral temporal lobe, and the bilateral Dorso-Lateral Pre-Frontal Cortex (DLPFC, Brodmann Area 46). The OFC and temporal lobes were intrinsically correlated with areas of mania-causing lesions. BA46 was intrinsically anti-correlated with mania-causing lesions. None of the control lesions were connected to these areas. Color Bar: Z-stat.

**Supplemental Figure 2:**
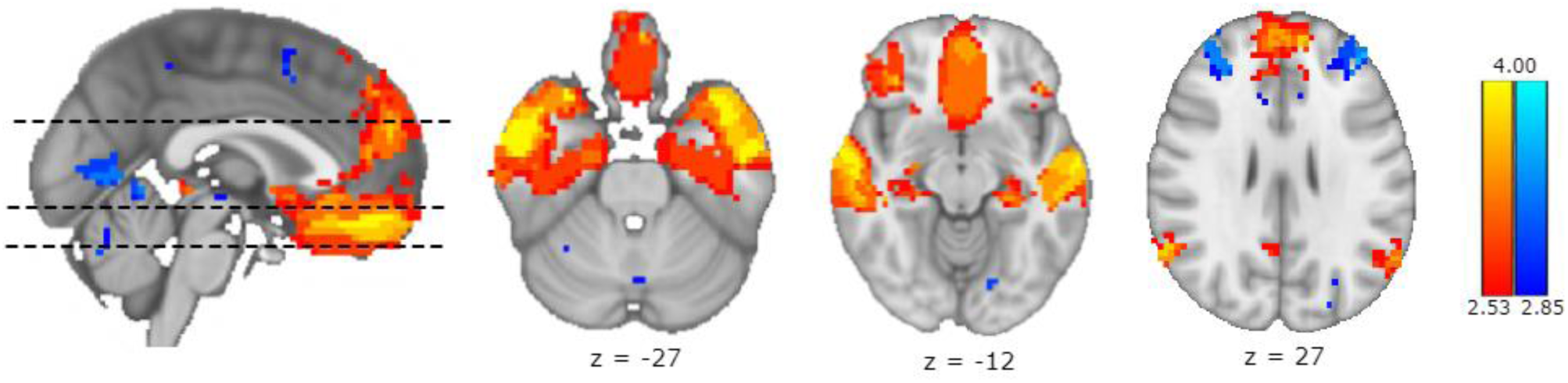
Mania Lesion Connectivity Compared to Size Matched Lesions Traced in a Single Plane: Network connectivity to mania causing lesions is compared to network connectivity in a set of control stroke lesions that have been selected on the basis of similarity in size to the mania causing lesions. A voxelwise Liebermeister test reveals regions significantly (p-FDR <.05) more functionally connected to mania-causing lesions than controls lesions. Regions of positively correlated activity to the ROI are shown in hot colors and negatively correlated activity in cold colors. Three regions of significant differences in connectivity were observed: The Orbitofrontal Cortex (OFC), the bilateral temporal lobe, and the bilateral Dorso-Lateral Pre-Frontal Cortex (DLPFC, Brodmann Area 46). The OFC and temporal lobes were intrinsically correlated with areas of mania-causing lesions. BA46 was intrinsically anti-correlated with mania-causing lesions. None of the control lesions were connected to these areas. Color Bar: Z-stat.

**Supplemental Figure 3:**
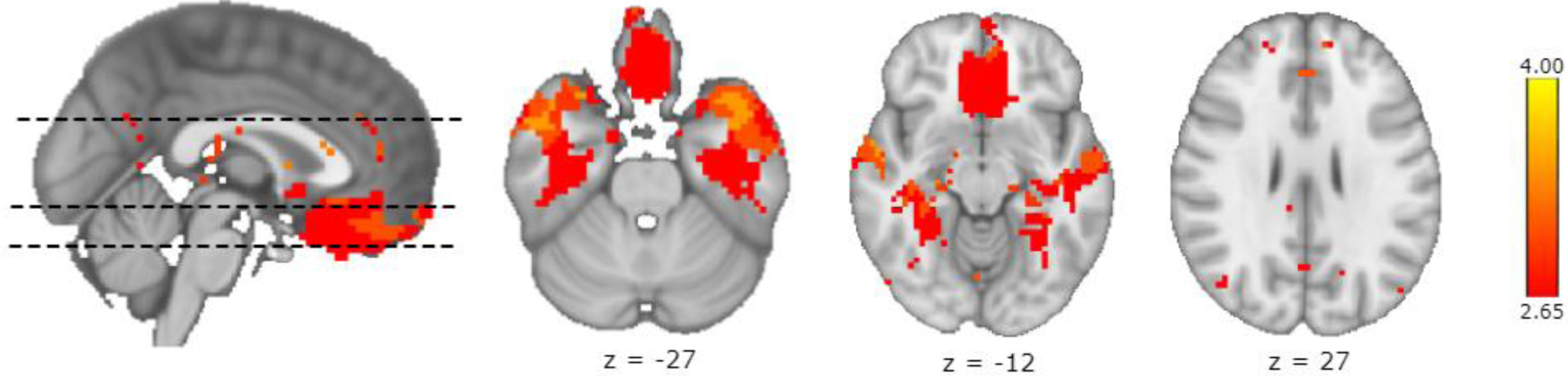
Mania-causing infarcts Alone Compared to Control Lesions. Network connectivity to mania causing lesions due to an ischemic infarct (n=13) are compared to network connectivity in a set of control stroke lesions (n=23). A voxelwise Liebermeister test reveals regions significantly (p-FDR <.05) more functionally connected to mania-causing lesions than controls lesions. Regions of positively correlated activity to the ROI are shown in hot colors. Two regions of significant differences in connectivity were observed: The Orbitofrontal Cortex (OFC), the bilateral temporal lobe. The OFC and temporal lobes were intrinsically correlated with areas of mania-causing lesions. None of the control lesions were connected to these areas. Color Bar: Z-stat.

**Supplemental Table 2:**
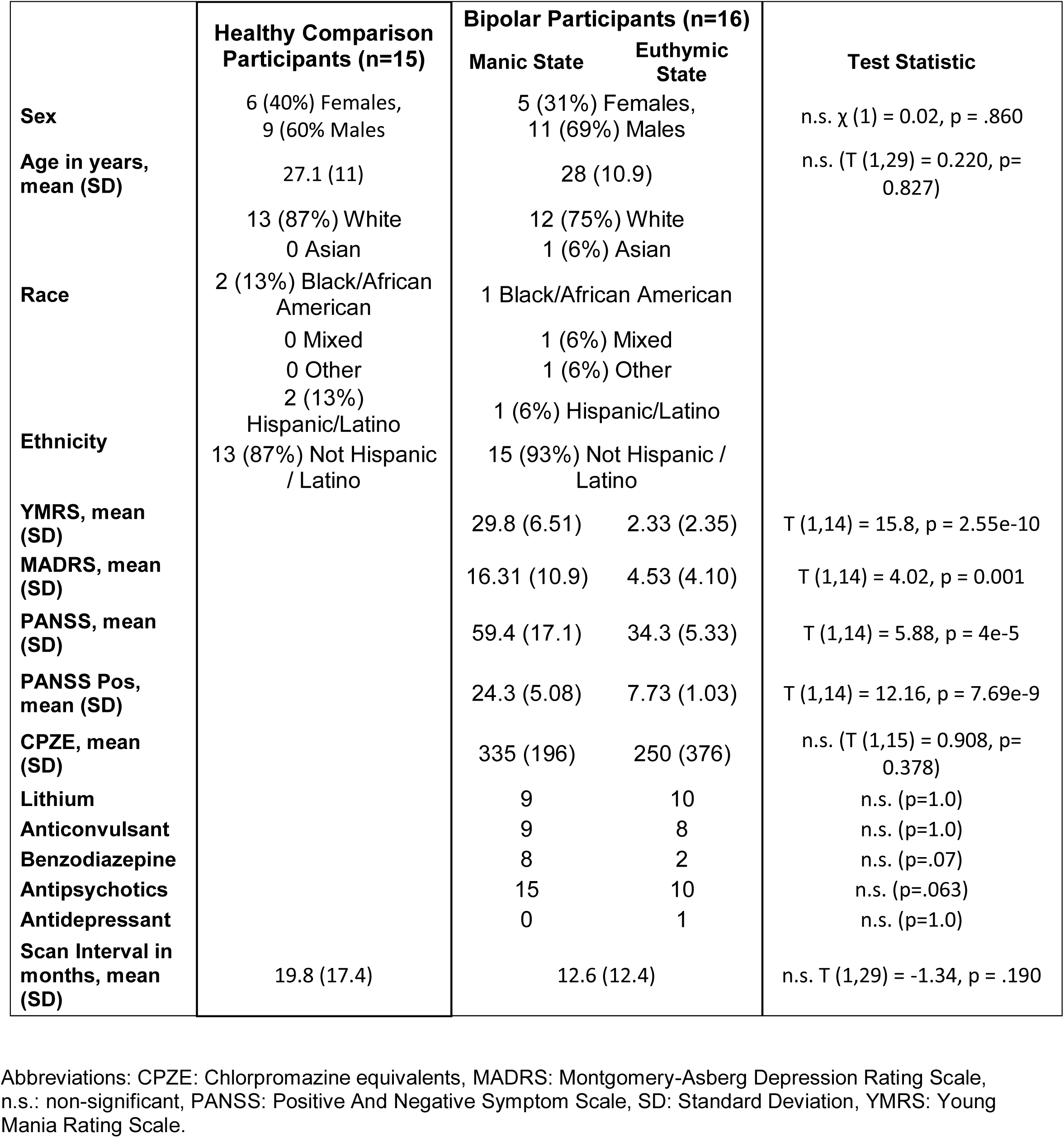
Longitudinal Participant Demographics and Clinical Characteristics.

### Bipolar Imaging Analysis Covarying for Medication Status

We did not observe any significant within-subject changes in medication regimen in the bipolar participants. There was a trend towards decreased presence of benzodiazepines and antipsychotics during the manic state. Although not significant (especially after multiple comparison correction), we re-performed the ROI-ROI analyses in the bipolar cohort examining functional connectivity in a mixed analysis with the presence or absence of these medications in regimen as a time-varying covariate. Mixed model covariance type was unstructured. Fixed effects included mood state. Random effects included medication status (prescribed or not prescribed) for benzodiazepines and for antipsychotics. BA46-amygdala connectivity continued to show a significant effect of mood state when covarying for (F(1,15)=38.9, p=.000016) Likewise, temporal-VLPFC connectivity also remained significant (F (1,16.1)=48.4, p=.000003).

### Bipolar Imaging Controlling for Psychiatric Comorbidity

We did not exclude participants on the basis of a substance use disorder so we repeated these ROI-ROI connectivity comparisons excluding any participant that met criteria for a substance use disorder at either time point. This analysis continued to demonstrate significant effects of mood with similar or stronger effect sizes: BA46-amygdala T(1,9)= 3.875, p=.004, Cohen’s d=-1.23. Mean zFC mania -.054, Mean zFC euthymia - .256.Temporal-VLPFC T(1,9)= 6.727 p=.000086, Cohen’s d=2.14. means zFC mania -.058, Mean zFC euthymia .288.

**Supplemental Figure 4:**
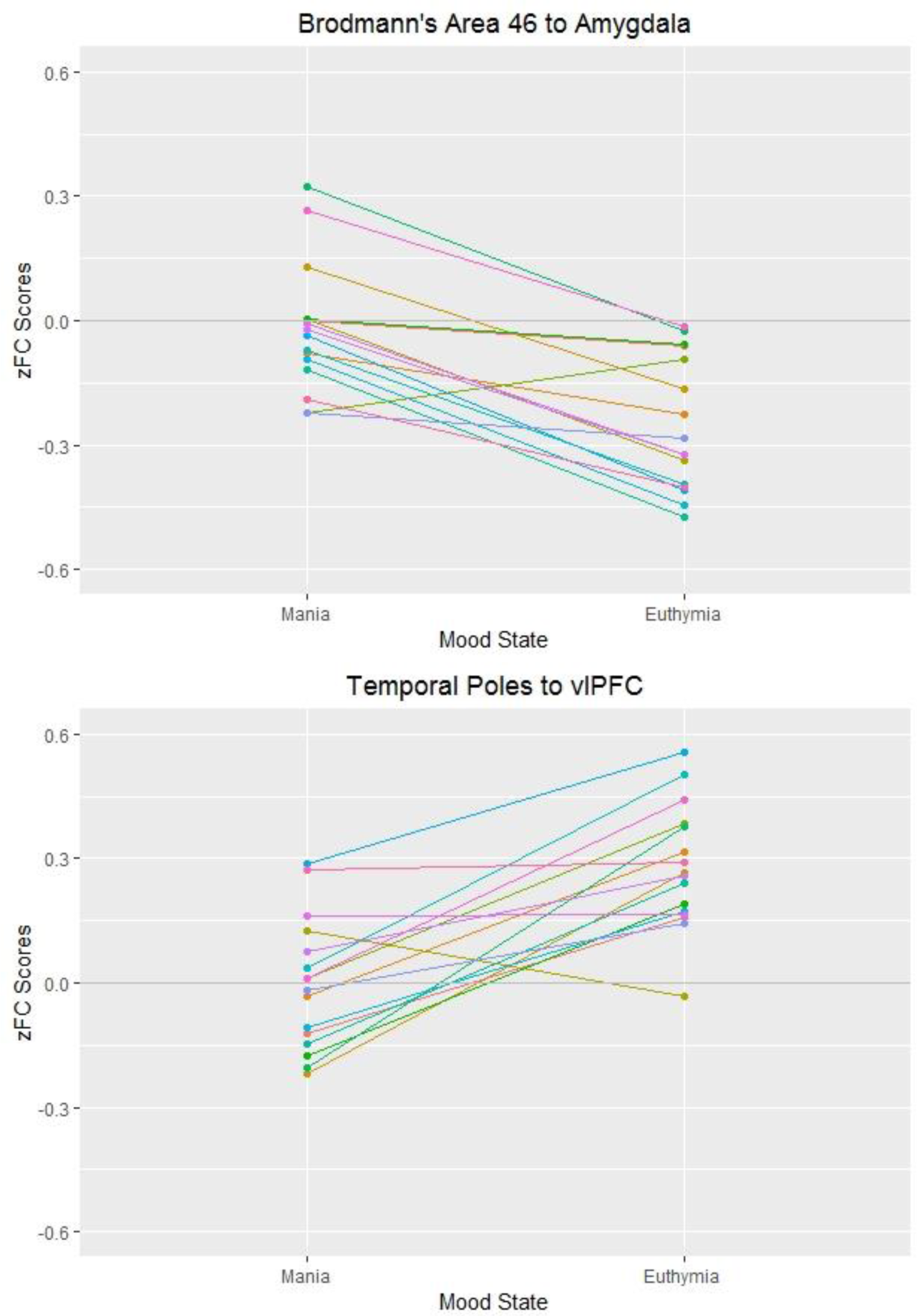
Individual Participant Connectivity Values in Mania and Euthymia. ROI to ROI connectivity values for each bipolar participant in each mood state. A) Individual connectivity trajectories between temporal lobes and right VLPFC in mania and euthymia. B) Individual connectivity trajectories between BA46 and left Amygdala in mania and euthymia.

**Supplemental Table 3:** Non-Longitudinal Bipolar Replication Participants:

|  | Demographics |  | Statistic |
| --- | --- | --- | --- |
|  | Bipolar-Manic<br>Subjects<br>(n=26) | Bipolar-Euthymic<br>Subjects<br>(n=21) | Mania vs<br>Euthymia |
| **Sex** | 20 Male, 6<br>Female | 13 Male, 8<br>Female | p=.263 |
| **Age in years,**<br>**mean (SD)** | 27.7 (10.5) | 33 (12.9) | p=.094 |
| **# of**<br>**hospitalizations,**<br>**mean (SD)** | 3.79 (4.19) | 4.61 (3.8) | p=.423 |
| **Duration of**<br>**illness in years,**<br>**mean (SD)** | 11.6 (12.1) | 7.32 (7) | p=.308 |
| **YMRS, mean**<br>**(SD)** | 28.6 (6.71) | 3.62 (3.84) | p= 4.75 x 10<sup>-9</sup> |
| **MADRS, mean**<br>**(SD)** | 14.3 (8.03) | 6.09 (5.48) | p = 2.54x10<sup>-4</sup> |
| **PANSS, mean**<br>**(SD)** | 56 (16.2) | 39 (8.85) | p=2.21 x 10<sup>-4</sup> |
| **PANSS positive**<br>**subscale, mean**<br>**(SD)** | 22.7 (5.35) | 9.9 (3.51) | p=2x10<sup>-8</sup> |
| **Anticonvulsants** | 8/26 | 10/21 | ns |
| **Antipsychotics** | 25/26 | 16/21 | p=.041 |
| **CPZ equivalents,**<br>**mean (SD)** | 396 (349) | 203 (239) | p=0.014 |
| **Benzodiazepines** | 8/26 | 4/21 | ns |
| **Lithium** | 18/26 | 8/21 | p=.033 |
| **Antidepressants** | 1/26 | 3/21 | ns |

Abbreviations: CPZE: Chlorpromazine equivalents, MADRS: Montgomery-Asberg Depression Rating Scale, n.s.: non-significant, PANSS: Positive And Negative Symptom Scale, SD: Standard Deviation, YMRS: Young Mania Rating Scale.

## Acknowledgments

This work was supported by the National Institutes of Health:

K23MH100623 (RB)

RO1MH78113 (MK)

K24MH104449 (DO)

R01MH109687 (MH)

